# Multiplex viral tropism assay in complex cell populations with single-cell resolution

**DOI:** 10.1101/2021.07.12.451549

**Authors:** Keng Choong Tat, Guo Ke, Daryl Lim Shern, Wei Leong Chew

## Abstract

Gene therapy constitutes one of the most promising modes of disease treatments. Two key properties for therapeutic delivery vectors are the transduction efficiency (how well the vector delivers therapeutic cargo to desired target cells) and specificity (how well it avoids off-target delivery into the other unintended cells within the body). Here we developed a novel technology that enables multiplex measurement of transduction efficiency and specificity, particularly by measuring how libraries of delivery vectors transduce libraries of diverse cell types. We demonstrated that pairing high-throughput measurement of AAV identity with high-resolution single-cell RNA transcriptomic sequencing maps how natural and engineered AAV variants transduce individual cells within human cerebral and ocular organoids. This library-on-library technology is important for determining the safety and efficacy of therapeutic delivery vectors.

## Introduction

Adeno-associated viruses (AAVs) are medically and commercially attractive gene delivery vectors due to the recent successes in FDA and EMA approvals for AAV-based gene therapies, as exemplified by Glybera for the treatment of lipoprotein lipase deficiency, Luxturna for the treatment of inherited retinal disease, and Zolgensma for the treatment of paediatric spinal muscular dystrophy^1,2,3,4,5^. The therapeutic applications of AAV span from targeting small tissues in the eye to systemic distribution throughout multiple organs including difficult-to-access systems such as the nervous system and vasculature^1^. This versatility is enabled by the ability to manipulate the AAV protein capsid sequence, which in turns changes the serotype and confers preferential tropism towards desired tissues. While considerable efforts have been devoted to identify optimal capsid proteins for successful therapy, early studies comparing the performance of different AAV serotypes are often of low-throughput and costly. A first limitation is that each cell line or animal is usually only transduced by a single AAV serotype and hence to evaluate multiple different serotypes would require a similar increase in independent replicates; this is in part because readouts employed for transduction efficiency assays tend to be non-multiplexable, such as quantification by immunohistology or fluorescence reporter proxy^6,7,8,9,10^, which means that each sample could only be treated by a single vector test candidate. A second limitation is that the sensitivity of transduction assays tends to require aggregation across many cells and vector copies, and hence the resolution is limited to the tissue level instead of the often required cellular level. Such single-plex approaches limit comparison to only a small handful of AAV serotypes in a similarly small number of target cells or tissues. In recent years, transduction assays of higher throughput have been devised by harnessing sequencing as a readout for transduction efficacy, whereby multiplex libraries of AAVs bearing nucleotide barcodes are administered to the target cells or tissues and the best-performing AAV serotype are identified by sequencing the nucleotide barcodes^11,12,13,14^. However, the techniques employed so far have been limited to bulk tissues, which do not offer the resolution needed to profile how efficiently or specifically each AAV serotype transduces specific subset(s) of cells within a complex tissue population^15,16,17,18^.

In this study, we developed a technology pipeline that enables multiplex measurement of AAV transduction efficiency and specificity for each cell type within a heterogeneous population. We barcoded AAV serotypes according to a new design principle, applied this AAV library on complex mixtures of cell types, conducted single-cell sequencing ^19,20,21,22,23,24^ to identify both the cell type and the AAV barcodes the single cell contains, and deconvoluted these data into matrices of AAV serotype versus human cell types. We applied this technology in human organoids, which can recapitulate certain structural and cellular complexity of the human brain and eye, and identified how efficiently and specifically each AAV serotype transduces individual cell types found within the organoids. This technology enables a more comprehensive interrogation of delivery vector biodistribution that will impact safety and efficacy profiles of the therapeutic product.

## Results

### Study Design

In this study, we aim to provide a new framework for assessing multiplex viral tropism in complex tissues in a high-throughput manner and at single-cell resolution. To accomplish this aim, we first generated panels of AAV serotypes where the AAV cargo is uniquely differentiable from each other. Specifically, individual packaging vectors of each AAV serotype each contains an eGFP transgene that is barcoded by a unique 8bp sequence at its 3’ end prior to the polyadenylation tail sequence (Supplementary Fig 1A). AAVs were produced from these barcoded packaging plasmids and the pooled AAVs were then used to transduce heterogeneous populations of cells within human ocular and cerebral organoids. Following transduction and cargo expression within infected cells, the organoids were then dissociated for single-cell sequencing, so as to identify the cell type and the AAV barcodes that infected the particular cell (Fig. 1A-D). Modifications made to the genome reference file and genome transcript file allowed for the AAV barcoded transcripts to be aligned and clustered together with the RNA transcriptomics data for the assignment and visualization of each AAV serotype transcript to individual cells in the Loupe Browser software at single-cell resolution (refer to materials and methods).

**Figure 1.**
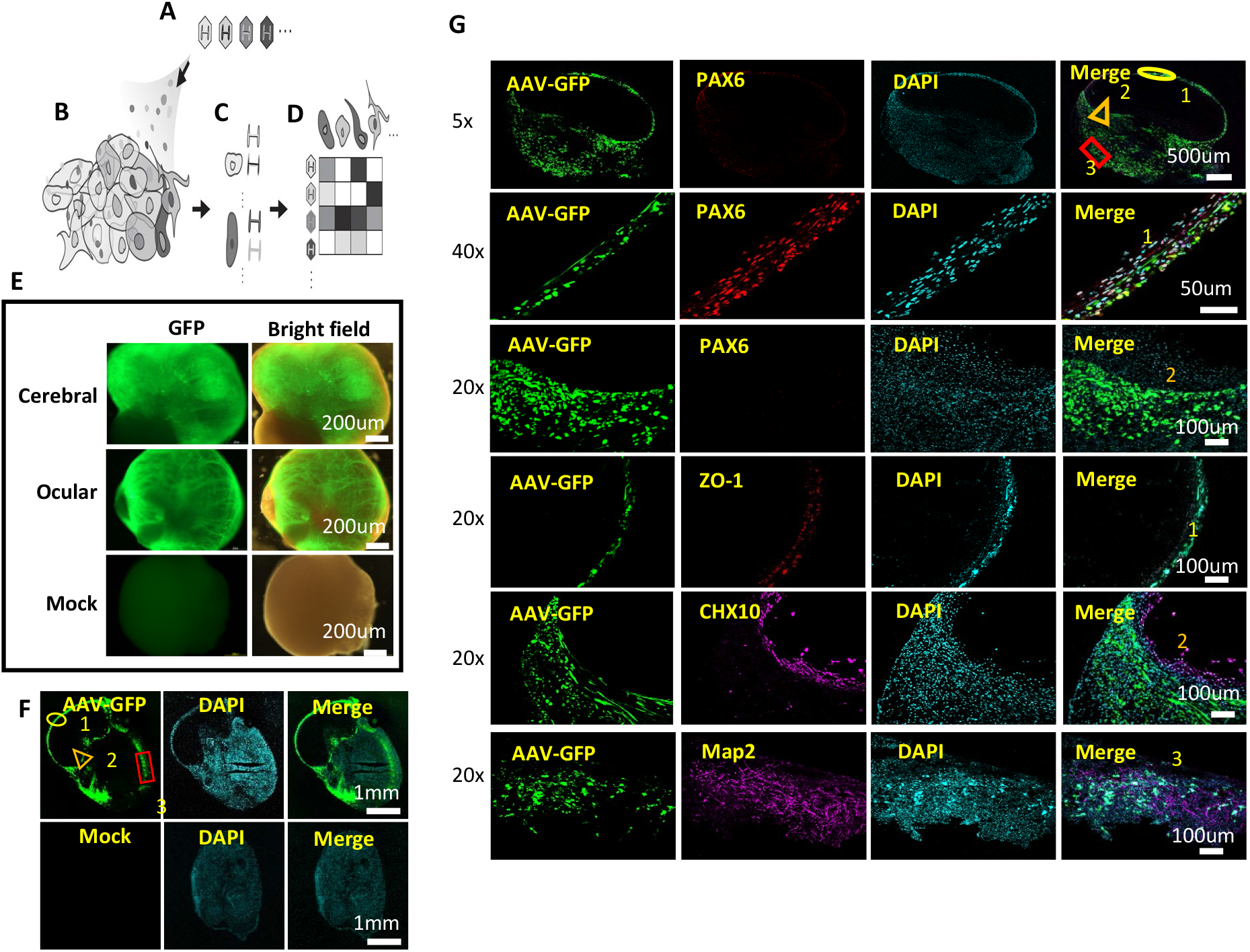
Framework for multiplex AAV tropism analysis at single cell resolution for pooled AAV transduced human ocular and cerebral organoids. (A) AAV serotype variants consisting a multitude of capsid serotypes are individually packaged with uniquely identifiable genomes with differentiable barcode sequences. (B) Library of AAV variants transduce a heterogenous population of cells (e.g. organs, tissues, organoids, admixtures). (C) In single-cell sequencing, nucleic acids (which can include RNA, DNA) within each cell are tagged by a unique cell-specific single-cell-sequencing nucleotide tag, then sequenced, and each cell is identified by its RNA transcriptome and/or DNA genome. (D) The matrix of which cell identity is being transduced by which AAV serotype can be created by matching the AAV variants with their respective transduced cells through the matching of their cell-specific single-cell-sequencing nucleotide tags. (E) Gross Morphology of cerebral and ocular organoids infected with AAV serotypes pool. Barcoded GFP-AAV-Pool (1 × 10^10^ vg/per serotype) expressing eGFP were used for transduction of cerebral and ocular organoids for 7 days. Low magnification microscopy showed GFP-positive signals in cells within most regions. *Mock* indicates negative control of un-transduced organoids. (F) Cross-section of AAV-infected ocular organoids showed GFP expression in different regions of the organoids indicating high transduction efficiency of the pooled AAV serotypes. Image inserts represent regions with predominant cell-types: *1* corneal cell-types, *2* retinal cell-types, *3* neuronal cell-types. (G) Immunofluorescence staining of cellular markers and GFP protein for identification of cell types transduced by the AAV serotypes pool. PAX6 - ocular epithelial or endothelial cells. CHX10 -specification and morphogenesis of the sensory retina. ZO-1-corneal endothelia marker. MAP2 –neuronal marker. DAPI marks nuclei.

### Barcoded AAVs transduce diverse tissue subtypes in human ocular and cerebral organoids

We reasoned that human ocular and cerebral organoids serve as models that represent the complexity of human tissues comprising multiple cellular subtypes. The organoids were cultured by differentiating H1 and H9 lineage of human ES cells on petri dishes for 6 weeks (Supplementary Fig. 2A), following which the ocular organoids were characterized by immuno-staining for common ocular tissue cellular markers S100β, PAX6, CHX10, RAX, CD31 and αSMA (Supplementary Fig. 2B) and the cerebral organoids were characterized by immuno-staining with common neural tissue cellular markers S100β, NeuN and Map2 (Supplementary Fig. 2C). Both the ocular and cerebral organoids express different cellular markers in distinct cellular layers, indicating heterogenous tissue subtypes within the organoids. The barcoded AAV pools (1 × 10^10^ vg/per serotype) were then administered upon the cerebral and ocular organoids. Culturing these organoids for a further 7 days resulted in strong GFP-positive signals in cells within most regions of the organoids indicating transduction and expression of the GFP cargo common among the pooled AAVs (Fig. 1E-F). Co-localization of eGFP with several different cellular markers also confirmed that the pooled AAVs transduced diverse tissue subtypes within the human ocular and cerebral organoids (Fig. 1G).

**Figure 2.**
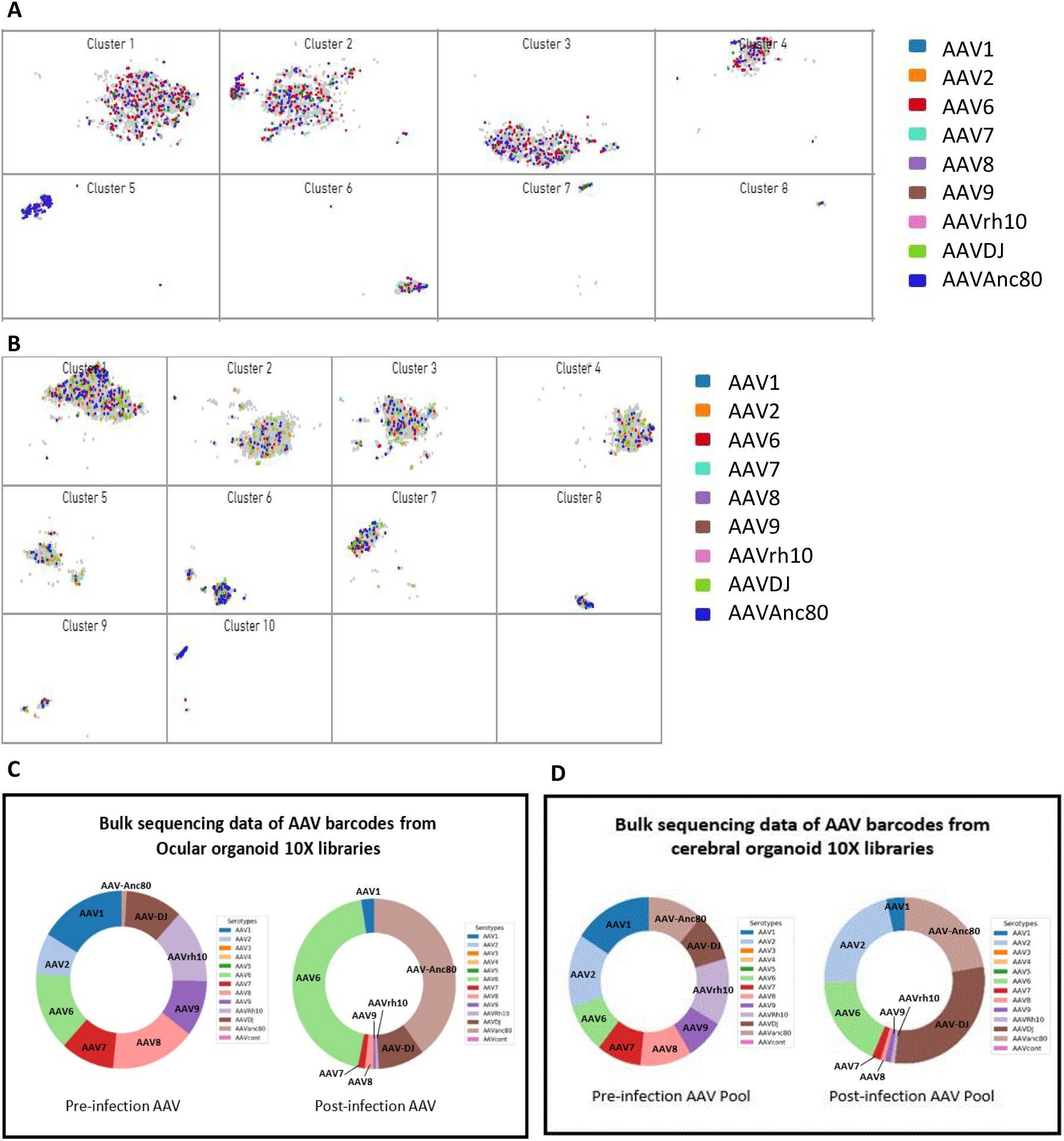
Single cell assignment of barcoded AAV transcripts for human ocular and cerebral organoids versus bulk analysis of AAV tropism. (A) t-SNE plots showing individual cells in ocular organoids transduced with different AAV serotypes (each serotype represented by one color), in each of the 8 clusters. Cluster 9 and 10 were omitted in this plot as no barcoded transcripts were assigned to these clusters. Plots with breakdown of each serotype for each cluster can be found in Supplementary Figure 4. (B) t-SNE plots showing individual cells in cerebral organoids transduced with different AAV serotypes (each serotype represented by one colour), in each of the 10 clusters. Plots with breakdown of each serotype for each cluster can be found in Supplementary Figure 5. (C) Bulk analysis of transduced ocular organoids by amplicon-sequencing on MiSeq sequencer. Results from bulk sequencing analysis is in agreement with the single-cell analysis plots processed with Cell Ranger pipeline, indicating that this assay enables accurate measurement of AAV tropism with single-cell resolution, beyond traditional bulk sequencing approach. (D) Bulk analysis of transduced cerebral organoids by amplicon-sequencing on MiSeq sequencer. Results from bulk sequencing analysis using custom Python script is in agreement with the single-cell analysis plots processed with Cell Ranger pipeline, indicating that this multiplex screening platform enables accurate measurement of AAV tropism with single-cell resolution, beyond traditional bulk sequencing approach.

### Single cell RNA transcriptomics clustering and assignment of AAV barcoded mRNA transcripts in transduced ocular and cerebral organoids at single cell resolution

After the human ocular organoids were transduced by the AAV libraries as described above, they were trypsinized into single cells as input for single-cell library preparation and sequencing (Materials and Methods). For ocular organoid, the transcriptomes of 5849 cells within the organoid (Sample number tested = 3) were profiled with mean reads per cell at 122688 and median genes per cell at 1022. Using K-means clustering, we were able to define 10 clusters of cells within the ocular organoids based on their transcriptomic profile (Supplementary Fig. 3A and Supplementary Data II). Each cluster of cells were uniquely identified by their top 10 expressing genes within the cluster (Supplementary Fig. 3B and Supplementary Data II). Fig. 2A shows a summary of all assignment of the AAV transcripts with each colour representing one AAV serotype, at single cell resolution. For more detailed analysis, the plot can be broken down to visualize individual serotype for each cluster (Supplementary Figure 4). To demonstrate that the methodology is easily applied on different complex tissues, we also conducted the same single-cell tropism assay on human cerebral organoids, which contain different populations of cell types compared to the ocular organoids. For the cerebral organoid, single-cell sequencing profiled the transcriptomes of 15466 cells within the organoid (Sample number tested = 2) with mean reads per cell at 23315 and median genes per cell at 902. Similarly, using K-means clustering, we were able to define 10 clusters of cells within the ocular organoids based on their transcriptomic profile (Supplementary Fig. 3C and Supplementary Data III). Each cluster of cells were uniquely identified by their top 10 expressing genes (Supplementary Fig. 3D and Supplementary Data III). Fig. 2B shows a summary of the assignment of all the AAV transcripts with each colour representing one AAV serotype, at single cell resolution. For more detailed analysis, the plot can be broken down to visualize individual serotype for each cluster (Supplementary Figure 5).

**Figure 3.**
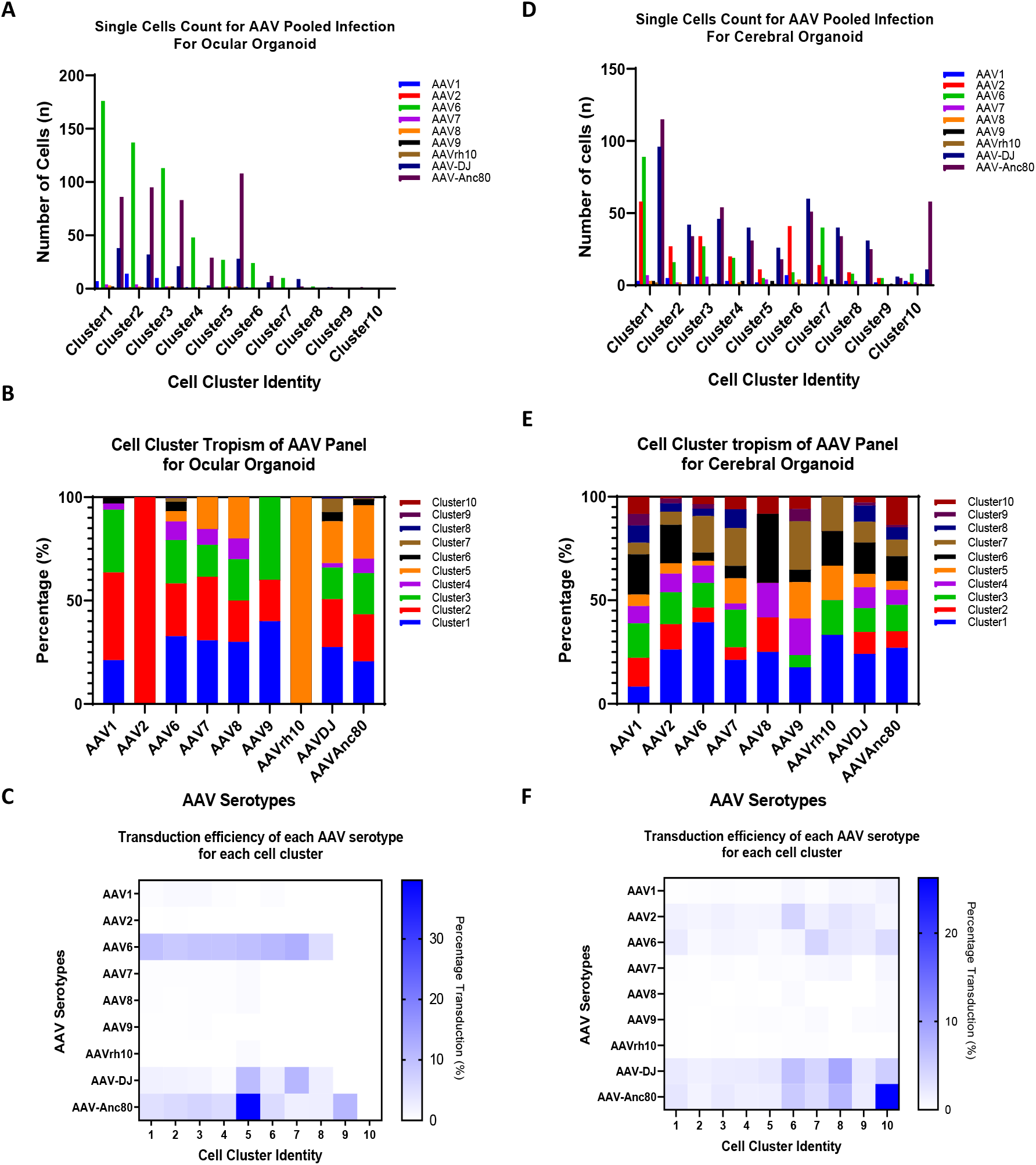
High-throughput AAV tropism measurement and analysis for human ocular and cerebral organoids. (A) Counts of cells that are transduced with each AAV serotype in each cluster. Data showed unique transduction level of each AAV serotype across the different cell clusters within human ocular organoids. (B) AAV cell cluster tropism in transduced human ocular organoid. Results demonstrated that the tropism of each AAV serotype varied across the different cell clusters and are distinct from other AAV serotypes. (C) The transduction efficiency of each AAV serotype for each cell cluster is visualized as the percentage of cells transduced in a heat map. Using this method, this assay enables identification of (i) the most efficient AAV serotype for each cell cluster and (ii) the most specific AAV serotype for the target cell type of choice (i.e. lowest transduction of other non-desired cell types). (D) Graph of counts of cells that are transduced with each AAV serotype in each cluster. Data showed unique transduction level of each AAV serotype across the different cell clusters within human cerebral organoids. (E) Graph of AAV cell cluster tropism in transduced human cerebral organoid. Results demonstrated that the tropism of each AAV serotype varied across the different cell clusters and are distinct from other AAV serotypes. (F) The transduction efficiency of each AAV serotype for each cell cluster is visualized as the percentage of cells transduced in a heat map plot. Using this method, this assay enables identification of (i) the most efficient AAV serotype for each cell cluster and (ii) the most specific AAV serotype for the target cell type of choice (i.e. lowest transduction of other non-desired cell types).

Next, we compared the multiplex tropism assessment technology to bulk sequencing of the GFP barcodes, which is a method commonly used by recent studies to examine AAV transduction in bulk tissues. Data from bulk sequencing of ocular organoids is in concordance with the single-cell sequencing data aggregated across cells (Table 3 and Fig. 2C), with AAV-Anc80, AAV6 and AAV-DJ being the top 3 AAV serotypes that most efficiently transduce the ocular organoids in bulk or in aggregate among single-cells. Similarly, the data from bulk sequencing of the cerebral organoids also aligns with the single cell sequencing data with AAV2, AAV6, AAV-DJ and AAV-Anc80 as the top 4 AAV serotypes that can most efficiently transduce the cerebral organoids (Table 4 and Fig 2D).

Importantly, by extracting the read counts of the different AAV serotypes transcripts in each cell cluster, we were able to visualize the absolute (Fig. 3A and 3D) and relative (Fig. 3B and 3E) transduction efficiency of each AAV serotype across heterogeneous cell types within the organoids. For ocular organoids, when normalized against GAPDH across all clusters, AAV-Anc80 is identified as the most efficient serotype for targeting cell clusters 5 representing retinal-like cell types (RDH5^hi^, MITF^hi^), while AAV6 and AAVDJ are the most efficient serotype transducing cell cluster 7 representing epithelium-like cell types (TP63^hi^, KRT5^hi^) and cluster 8 representing neural stem-like cell types (PAX6^hi^, SOX2^hi^, MAP2^hi^) (Fig. 3C). Similarly, for the cerebral organoid, when normalized against GAPDH across all cell clusters, AAV2, 6 and Anc-80 were identified as serotypes that can efficiently transduce cluster 6 representing brain meningeal-like cells (DCN^hi^, SOX2^hi^, PAX2^hi^) while AAV6 and AAVDJ most efficiently transduce cluster 7 representing midbrain dopaminergic-like cells (RSPO2^hi^, SOX2^hi^, PAX6^hi^). In addition, the result suggests that AAVDJ is the most efficient serotype for cluster 8 representing astroglia or Schwann-like cell types (S100B^hi^) and AAV-Anc-80 is the most efficient serotype for cluster 10 representing microglial-like cells (UCP2^hi^) (Fig. 3F). These results show that the single-cell AAV tropism assay identifies different AAV serotypes with preferential tropism towards each subset of human cell types within the ocular or cerebral organoids.

## Discussions

To date, most of the published AAV tropism assays utilize low-resolution methods to perform relative comparison between a few AAV serotypes, conducted either *in vitro* using homogenous cell lines or *in vivo* using bulk tissue organs. In this study, we demonstrated a pipeline that enable high-throughput multiplexing of AAV libraries for relative comparison of transduction efficacies at single-cell resolution. We evaluated the tropism of a library of AAV serotypes consisting of natural (AAV1, 2, 6, 7, 8, 9, and rh10) and engineered AAVs (DJ and Anc-80) for their transduction efficacy across different single-cell niches within the same tissue organoid simultaneously. High-resolution quantification of every AAV serotype mRNA transcripts that are present in each single cell reveals the AAV serotype(s) that has preferential tropism towards individual cell types. Although the current demonstrated data employs the use of only 9 serotype variants, the assay could likely support substantially more variants as the barcoding strategy allows for simple scaling up (i.e., the current 8-nt barcoding can support 65K unique barcodes and serotypes, before implementation of error-tolerating or error-correcting encoding). The assay can also be applied beyond ocular or cerebral organoids to any tissue in culture or *in vivo*, especially when the targeted cellular subtypes have established cell type markers to facilitate annotation. This method can potentially be employed for clinical development by refining the selection of AAV serotypes for precise gene delivery to diseased tissues.

## Materials and Methods

### Organoids culture and condition

Briefly, the cerebral and ocular organoids were cultured in mTeSR1 medium (Stem Cell Technologies, cat. no. 85850). Human ES cell (H1 WA01 and H9 WA09) were treated by accutase to generate single cells. Then, 4000 cells were plated in each well of a V-bottom 96-well plate (Sematec Pte Ltd Code: 1009985) with low concentrations of basic fibroblast growth factor (bFGF 4 ng/ml) and 20 uM/ml Rho-associated protein kinase (ROCK) inhibitor (Y27632 Stem Cell). The next day, Embryonic Bodies (EB) were transferred into in low-attachment 96 well U-bottom plate with hESC medium (For 500 ml of medium, combine 400 ml of DMEM-F12, 100 ml of KOSR, 15 ml of ESC-quality FBS, 5 ml of GlutaMAX, 5 ml of MEM-NEAA and 3.5 μl of 2-mercaptoethanol) for cerebral organoid, Differentiation Medium DM (DMEM/F12, 4% knockout serum replacement (KOSR), 4% fetal bovine serum (ESC-quality FBS), 1× non-essential amino acids (NEAA), 1× Glutamax, 1× Pen-Strep. Filter it using a vacuum-driven 0.2-μm filter unit) for ocular organoid. EB were fed every other day for 6 days and then changed into neural induction media for cerebral organoid and into retinal differentiation medium (RDM: DM + 2% B27) for ocular organoid for the next 4 days. After the EB undergone neuro-ectodermal differentiation, they are transferred to Matrigel (Growth factor–reduced Matrigel, Bio-Lab 354230). Making matrigel in 1:1 dilution with cerebral organoid differentiation medium or corneal differentiation medium (CDM). 50ul of matrigel is added to each well and incubated for 30min in a 37°C incubator, followed by adding 100ul cerebral organoid differentiation medium with B27 (-) Vitamin A to each well and cultured for 48h. After 2-3 days, the aggregates (organoids) were transferred to 6-well clear flat-bottom ultra-low attachment plates. After 4 days of static culture with cerebral organoid differentiation medium with B27 (-) Vitamin A, the embedded organoids were transferred to an orbital shaker at 80 rpm within37°C, 5% CO2 incubator for long-term culture with cerebral organoid differentiation medium with B27 (+) Vitamin A.

### AAV plasmid cloning and virus production

The barcoded eGFP plasmids were constructed by introducing a short sequence TAATAAATCGATCGNNNNNNNN after the eGFP transgene stop codon in the plasmid backbone pZac2.1-CMV-eGFP.rgb, a gift from Luk Vandenberghe. Primers with overhanging barcode were designed for first round PCR to generate barcoded eGFP fragments that terminates at ITR sequences. A second round of nested PCR amplify shorter fragments of barcoded eGFP which are digested with restriction enzyme NheI and BamHI. Digested fragments are ligated with the vector backbone which is digested using the same restriction enzymes. The sequences of the clones were checked by Sanger sequencing. The representing barcodes for each AAV serotype are shown in Table 1. The serotype-specific pAAV-RepCap plasmids were constructed by cloning in the Cap genes from the different serotypes into the pAAV-RepCap backbone using Gibson assembly. The different serotypes Cap genes were ordered as gene blocks (IDT) and cloned into HindIII/PmeI-digested pAAV-RepCap backbone via Gibson assembly to construct the pAAV-RepCap with the different serotypes Cap genes. AAV viruses from different serotypes each bearing its own barcode were produced as per standard protocol^25^. Briefly, AAV were packaged via a triple transfection of 293AAV cell line (Cell Biolabs AAV-100) that were plated in a HYPERFlask ‘M’ (Corning) in growth media consisting of DMEM+glutaMax+pyruvate+10%FBS (Thermo Fisher), supplemented with 1X MEM non-essential amino acids (Gibco). Confluency at transfection was between 70–90%. Media was replaced with fresh pre-warmed growth media before transfection. For each HYPERFlask ‘M’, 200 μg of pHelper (Cell Biolabs), 100 μg of pRepCap [encoding capsid proteins for different serotypes], and 100 μg of pZac-CASI-GFP (barcoded) were mixed in 5 ml of DMEM, and 2 mg of PEI “MAX” (Polysciences) (40 kDa, 1 mg/ml in H_2_O, pH 7.1) added for PEI: DNA mass ratio of 5:1. The mixture was incubated for 15 min, and transferred drop-wise to the cell media. The day after transfection, media was changed to DMEM+glutamax+pyruvate+2%FBS. Cells were harvested 48–72 hrs after transfection by scrapping or dissociation with 1×PBS (pH7.2) + 5 mM EDTA, and pelleted at 1500 g for 12 min. Cell pellets were resuspended in 1–5 ml of lysis buffer (Tris HCl pH 7.5 + 2 mM MgCl + 150 mM NaCl), and freeze-thawed 3× between dry-ice-ethanol bath and 37 °C water bath. Cell debris was clarified via 4000 g for 5 min, and the supernatant collected. The collected supernatant was treated with 50 U/ml of Benzonase (Sigma-Aldrich) and 1 U/ml of RNase cocktail (Invitrogen) for 30 min at 37 °C to remove unpackaged nucleic acids. After incubation, the lysate was loaded on top of a discontinuous density gradient consisting of 6 ml each of 15%, 25%, 40%, 60% Optiprep (Sigma-Aldrich) in an 29.9 ml Optiseal polypropylene tube (Beckman-Coulter). The tubes were ultra-centrifuged at 54000 rpm, at 18 °C, for 1.5 hr, on a Type 70 Ti rotor. The 40% fraction was extracted, and dialyzed with 1×PBS (pH 7.2) supplemented with 35 mM NaCl, using Amicon Ultra-15 (100 kDa MWCO) (Millipore). The titer of the purified AAV vector stocks were determined using real-time qPCR with ITR-sequence-specific primers and probe^26^, referenced against the ATCC reference standard material 8 (ATCC).

### In vitro transduction of organoids

AAV serotypes pool was created by pooling each AAV serotype at 1 × 10^10^ vg, giving a final viral copy of 9 × 10^10^ that is used for the transduction of organoids in each well of a 24-well plate. AAV1, 2, 6, 7, 8, 9, rh10, DJ and Anc80 serotypes were used for the pooling. Organoids were transduced for 7-10 days before harvesting for sequencing, fluorescence imaging, and histochemistry.

### Immunofluorescence histochemistry

Organoids were fixed in 4% paraformaldehyde for 4hrs at 4°C followed by washing in PBS three times for 15 min. Organoids were allowed to sink in 30% sucrose overnight and then embedded in OCT and cryosectioned at 12um. Sections were permeabilized in 0.2% Triton X-100 in PBS and blocked in block buffer (2% BSA 5% fetal bovine serum) for 1 h at room temperature. Sections were subsequently incubated with the indicated primary antibodies at a 1:100 dilution in block buffer at 4°C overnight. Secondary antibodies used were donkey Alexa Fluor 488, 568 and 647 conjugates (Invitrogen, 1:1000). After staining with 4′,6-diamidino-2-phenylindole (DAPI) (Sigma-Aldrich) in PBS for 5 min, slides were mounted in Vectashield anti-fade reagent (Vector Laboratories). Confocal imaging was performed with Leica TCS SP8 DLS LightSheet microscope. Primary antibodies: PAX6 (rabbit, abcam ab5790), CHX10 (rabbit, abcam ab133636), ZO-1 (mouse, Thermofisher ZO1-1A12), MAP2 (chicken, abcam ab5392). S100β (rabbit, abcam ab52642), RAX (rabbit, abcam ab23340), CD31 (mouse, abcam ab23340), aSMA (rabbit, abcam ab5694), DAPI (49,6-diamidino-2-phenylindole). NeuN (mouse, Sigma-Aldrich MAB377).

### Amplicon barcode sequencing and analysis

Transduced organoid samples were harvested as single cells and processed through the 10X Chromium machine for cell barcoding of the transcripts. The total cDNA was purified via the 10X workflow and 5ul was aliquoted for custom bulk-sequencing. The rest of the cDNA were used to proceed with the remaining 10X workflow for single-cell sequencing. Custom primers were designed for a first round of 20-cycle PCR of the target site containing the AAV barcodes as shown in Table 1. Target bands are extracted using gel extraction and a second round of 15-cycle PCR were used for adding P5 and P7 adapter sequences to the enriched fragments and the final libraries were cleaned up by gel extraction. Primers used for library construction are shown in Table 2. Library concentrations were determined using a Qubit dsDNA HS kit (Agilent). NGS sequencing were carried out on the MiSeq using 2×75 bp PE run with 20% PhiX spike-in. An in-house python script was utilized to search for the 8 unique nucleotide barcode sequences representing each serotype within the MiSeq FASTQs generated from the MiSeq run of the amplicon libraries and the total count was tabulated for each barcode sequence for each sample (Refer to PythonScriptforBulkAnalysis package).

### Single cell sequencing and RNA transcriptomic analysis

Samples were prepared as indicated in the 10X Genomics Single Cell 3′ v2 Reagent Kit user guide. The single-cell libraries were prepared by following the manufacturers’ protocol followed by sequencing on an Illumina HiSeq4000 flow cell. The sequencing data were processed by the standard Cell Ranger pipeline using the modified gtf and genome manifest files. Briefly, the samples were washed twice in PBS (Life Technologies) + 0.04% BSA (Sigma) and re-suspended in the same solution. Sample viability was assessed using Trypan Blue (Thermo Fisher) under a light microscope. Following viability counting, the appropriate volume for each sample was calculated for a target capture of 10,000 cells and loaded onto the 10x Genomics single-cell-A chip along with other reagents and barcoded beads by following the protocol guide. The chip is then loaded onto a 10X Chromium machine for droplet generation and samples were transferred onto a pre-chilled strip tube (Eppendorf), and reverse transcription was performed using a 96-well thermal cycler (Thermo Fisher). After the reverse transcription, cDNA was recovered using Recovery Agent provided by 10X Genomics, followed by Silane DynaBead clean-up (10X Genomics). Purified cDNA was amplified for 12 cycles before being cleaned up using SPRI-select beads (Beckman). Samples were diluted 4 times in water and ran on a Bioanalyzer (Agilent Technologies) to determine cDNA concentration. cDNA libraries were then prepared following the Single Cell 3′ Reagent Kits v2 user guide with appropriate PCR cycles based on the cDNA concentration as determined by the bioanalyzer. The molarity of the single cell libraries was calculated based on their library sizes as measured using a bioanalyzer (Agilent Technologies) and using the KAPA qPCR quantification (KAPA) method on a qPCR cycler (Roche). Samples were normalized to 10nM before sequencing. Each organoid sample was sequenced on a full lane on a HiSeq 4000 with the following run parameters: Read 1 - 26 cycles, read 2 - 98 cycles, index 1 - 8 cycles. Using the FASTQ files from each sample, the standard Cell Ranger Count command pipeline was performed for transcripts read alignment, UMI counting, and clustering (Amazon Web Services via the Ronin cloud platform). Raw data were processed using standard Cell Ranger transcriptomics command, while using modified genome reference file and the modified gtf file. For command lines for the modification of gtf file and genome reference file to include the barcoded GFP sequences, refer to Supplementary Data 4. Command lines for cell count using the modified files are also shown in Supplementary Data 4. Finally, upon successful cellranger count run, the output file should contain a cloupe.cloupe file which allow the the single-cell clusters and transcript counts to be visualized in the Loupe Browser software user interface (10X Genomics).

### Single Cell AAV tropism analysis

For the purpose of parallel sequencing of the AAV barcodes in single cells along with the RNA transcripts, the human genome reference file and the genome transcript file (gtf) were modified (Supplementary Data 4). Briefly, the names and barcodes of each AAV serotypes are manually included into both files that will be used for the execution of the Cell Ranger Count command pipeline in order to include the AAV barcode transcripts into the read alignment, UMI counting, and clustering. To include the AAV barcode representation in the genome reference file, the command line “>GFP1 TAAATCGATCGNNNNNNNN” is included for each barcode, where the 8Ns represent a unique 8 nucleotide barcode sequence. The command line “GFP me exon 1 19 - + - gene_id “GFP1”; transcript_id “GFP1”” was included in the genome transcript file for each AAV barcode representation added to the genome reference file (Supplementary Data 4). In the Loupe Browser, K-means based clustering were selected to define niche cell population within each type of organoid. The AAV barcoded transcripts can be visualized under Gene/Feature Expression Analysis. The number of cells that are transduced by each serotype in each cell niche are then visualized using the Cell Loupe software and counted, and further tropism analysis (Fig. 3) was conducted in GraphPad Prism. To determine the transduction efficiency of a specific viral vector against a specific cell niche, we calculated the percentage of cells of the specific cell niche which have been detected positive for the presence of the specific viral vector. To calculate the transduction efficiency of a specific viral vector against a specific cell niche, we calculated the frequencies with which the presence of the specific viral vector is detected in the cells of the specific cell niche, against the frequencies with which the presence of another viral vector is detected in the cells of the same specific cell niche. To determine the transduction specificity of a specific viral vector against a specific cell niche relative to other cell niches, we calculated the frequencies with which the presence of that specific viral vector is detected in the cells of the specific cell niche, against the frequencies with which the presence of the same specific viral vector is detected in the cells of other specific cell niches.

## Data availability

High-throughput sequencing data for both the bulk sequencing and single cell sequencing can be accessed via NCBI Sequence Read Archive database with SRA accession PRJNA742883 and BioProject accession code [PRJNA742883](http://www.ncbi.nlm.nih.gov/bioproject/742883). The in-house Python package can be found at the same accession ID.

## Acknowledgements

We would like to thank Nicholas Ong for providing the python script for the analysis of AAV barcodes for the MiSeq bulk sequencing data, Hong-Ting Prekop for advice on AWS cloud platform set-up for the single-cell data processing work and Lavina Sierra Tay for proof-reading the manuscript. This work is supported by Agency for Science, Technology and Research (A*STAR) and Industrial Alignment Fund Pre-Positioning (IAF-PP) grant H17/01/a0/012.

## Authors Contributions

C.T.K. and W.L.C. contributed to the conception of single-cell tropism technology framework. C.T.K., K.G. and W.L.C. contributed to the conception of organoids as model of complex tissue for single-cell tropism. C.T.K. and D.L. contributed to the conception of the design and generation of the barcoded AAVs. C.T.K contributed to the cloning of the barcoded AAV transfer plasmids and bioinformatics analysis of the sequencing data. C.T.K and D.L. contributed to the production of the barcoded AAVs. K.G. contributed to the culture and maintenance of the human organoids. C.T.K and K.G contributed to the transduction and single-cell library preparation and sequencing of the organoids. C.T.K and W.L.C contributed to the writing of the manuscript.

## Supplementary Data I

**Table 1.**

| Plasmids | Barcode | For barcoding |
| --- | --- | --- |
| pZac2.1-CMV-eGFP_A701 | ATCACGAC | AAV1 |
| pZac2.1-CMV-eGFP_A702 | ACAGTGGT | AAV2 |
| pZac2.1-CMV-eGFP_A706 | AACCCCTC | AAV6 |
| pZac2.1-CMV-eGFP_A707 | CCCAACCT | AAV7 |
| pZac2.1-CMV-eGFP_A708 | CACCACAC | AAV8 |
| pZac2.1-CMV-eGFP_A709 | GAAACCCA | AAV9 |
| pZac2.1-CMV-eGFP_A710 | TGTGACCA | AAV-rh10 |
| pZac2.1-CMV-eGFP_A711 | AGGGTCAA | AAV-DJ |
| pZac2.1-CMV-eGFP_A712 | AGGAGTGG | AAV-Anc80 |

**Table 2.**
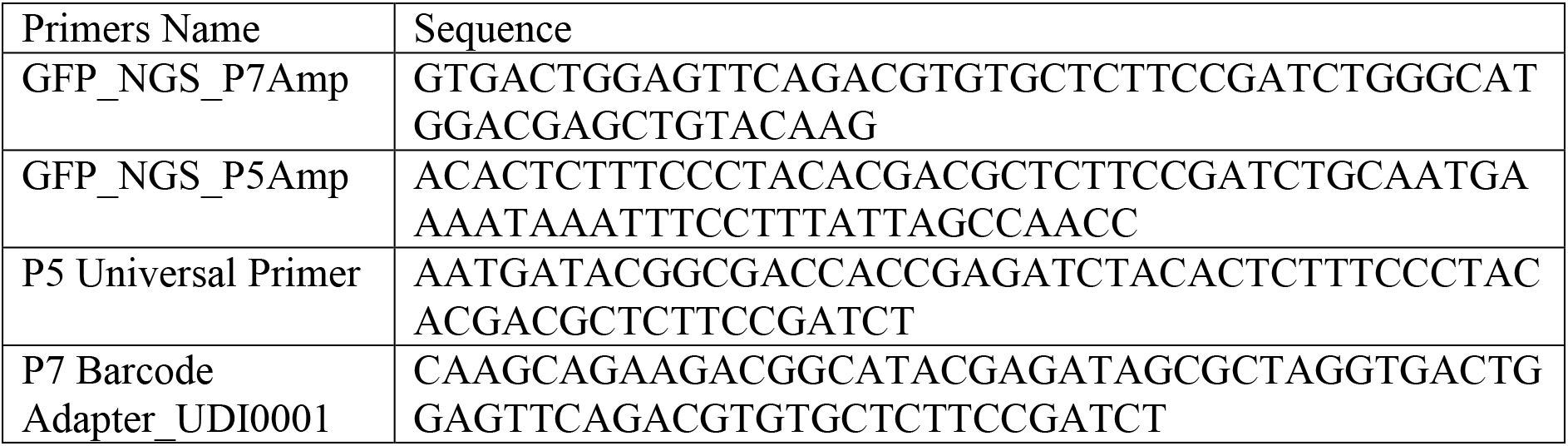

**Table 3.**

| AAV Barcodes | Bulk sequencing counts<br>for ocular organoid (%) | Single cell sequencing counts<br>for ocular organoid (%) |
| --- | --- | --- |
| A701 (AAV1) | 2.55 | 2.85 |
| A702 (AAV2) | 0.19 | 0.09 |
| A706 (AAV6) | 44.02 | 46.45 |
| A707 (AAV7) | 1.40 | 1.12 |
| A708 (AAV8) | 1.65 | 0.87 |
| A709 (AAV9) | 0.49 | 0.43 |
| A710 (AAVrh10) | 0.58 | 0.17 |
| A711 (AAVDJ) | 9.21 | 11.93 |
| A712 (AAVAnc80) | 39.88 | 36.07 |

**Table 4.**

| AAV Barcodes | Bulk sequencing counts<br>for cerebral organoid (%) | Single cell sequencing counts<br>for cerebral organoid (%) |
| --- | --- | --- |
| A701 (AAV1) | 3.63 | 2.62 |
| A702 (AAV2) | 21.72 | 16.08 |
| A706 (AAV6) | 17.88 | 16.45 |
| A707 (AAV7) | 1.76 | 2.40 |
| A708 (AAV8) | 0.91 | 0.87 |
| A709 (AAV9) | 1.17 | 1.24 |
| A710 (AAVrh10) | 0.80 | 0.44 |
| A711 (AAVDJ) | 29.60 | 28.97 |
| A712 (AAVAnc80) | 22.33 | 30.93 |

### Supplementary Figures

**Supplementary Figure 1.**
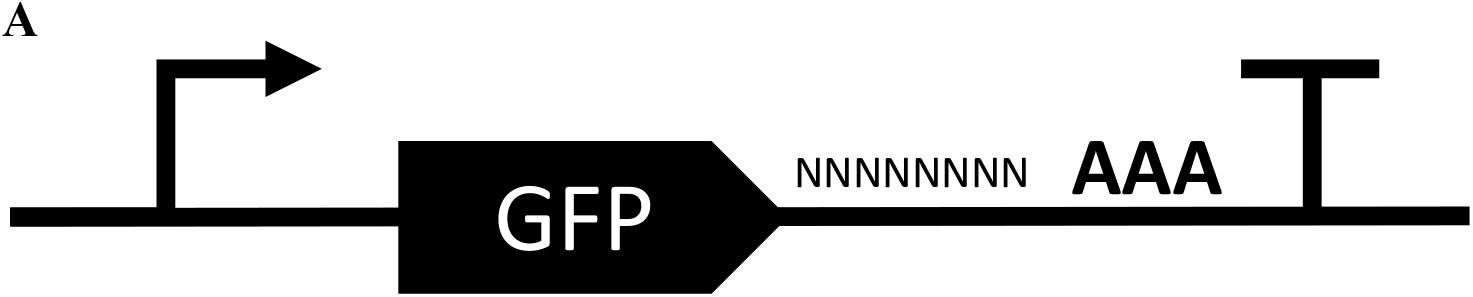
Design of AAV genomic cargo sequence for serotype barcoding and RNA transcripts capture for the modified 10X Cell Ranger pipeline for high-throughput single-cell analysis of AAV tropism. (A) Schematic of design of AAV genomic cargo for capture and analysis of serotype barcodes. A mammalian promoter is selected for expression of a non-host protein in the human organoid cells. An eGFP transgene with barcode is expressed and can be distinguished from host gene transcripts. A unique 8 base-pair barcodes is included after the stop codon and before the polyadenylation tail, designed to be within the 98 bases from captured tail for Cell Ranger analysis. A polyadenylation tail sequence is included for capture of RNA transcripts to the probes on 10X beads.

**Supplementary Figure 2.**
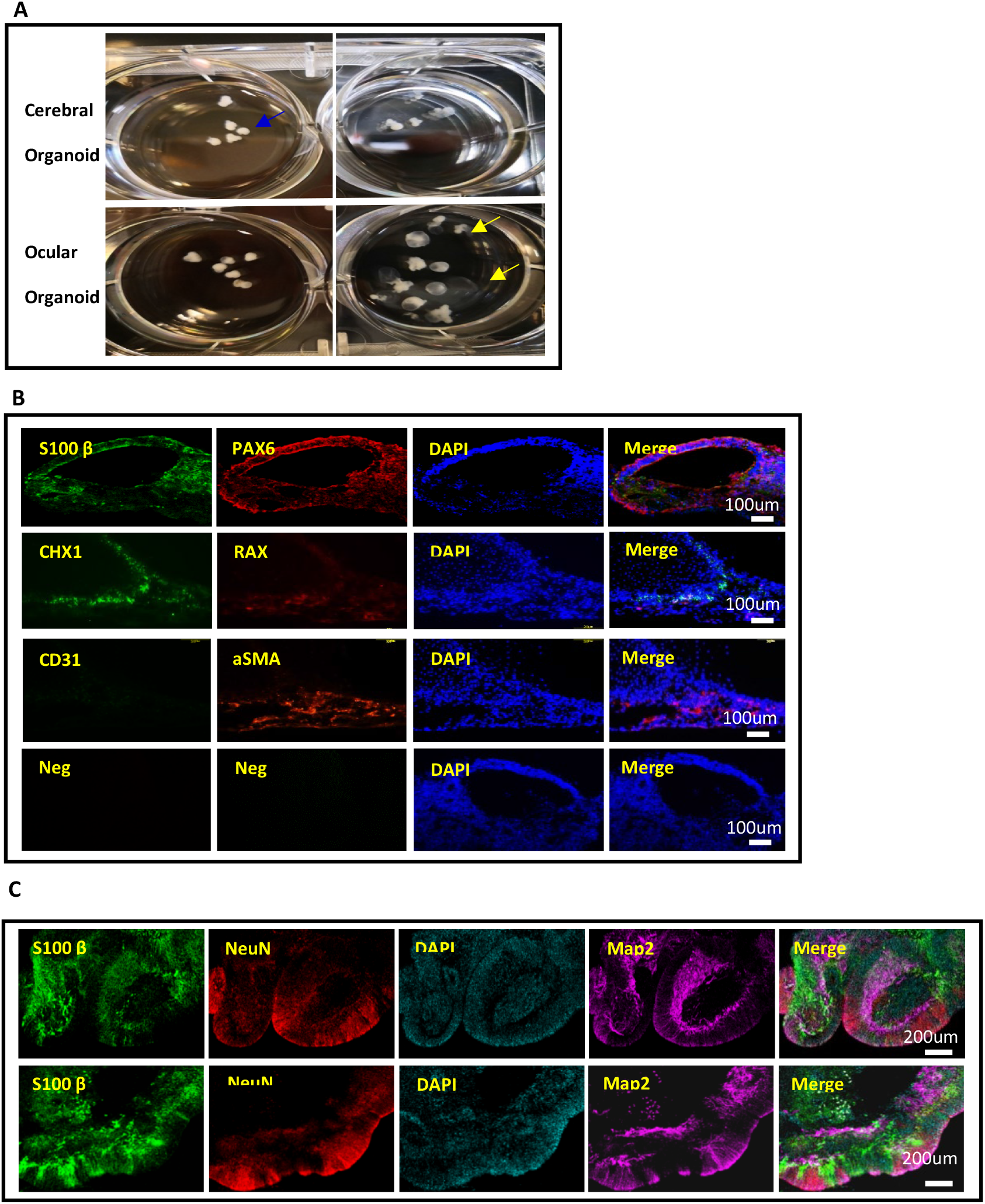
Ocular and cerebral organoids culture and characterization. (A) Gross morphology of developing human cerebral and ocular organoids cultured for 6 weeks. Low magnification bright-field images revealed fluid-filled cavities of ocular (yellow arrow) and solid brain (blue arrow) organoids. (B) Histology sections of ocular organoids were stained for cellular markers for cell-type characterization. S100β – neuronal crest and developed ocular. PAX6 – ocular epithelial or endothelial cells. CHX10 – specification and morphogenesis of the sensory retina. RAX – developing eye and initial specification of retinal cells. CD31 - Schlemm’s canal endothelial. aSMA - trabecular meshwork and stroma. DAPI (49, 6-diamidino-2-phenylindole) stain for nuclei. Neg – negative control. (C) Histology sections of cerebral organoids sections were stained with for cellular markers for cell-type characterization. MAP2 – Positive in all neural cells. NeuN – Neuronal marker. S100 β – detect brain proteins and express in the neuronal cells. DAPI- stain for nuclei.

**Supplementary Figure 3.**
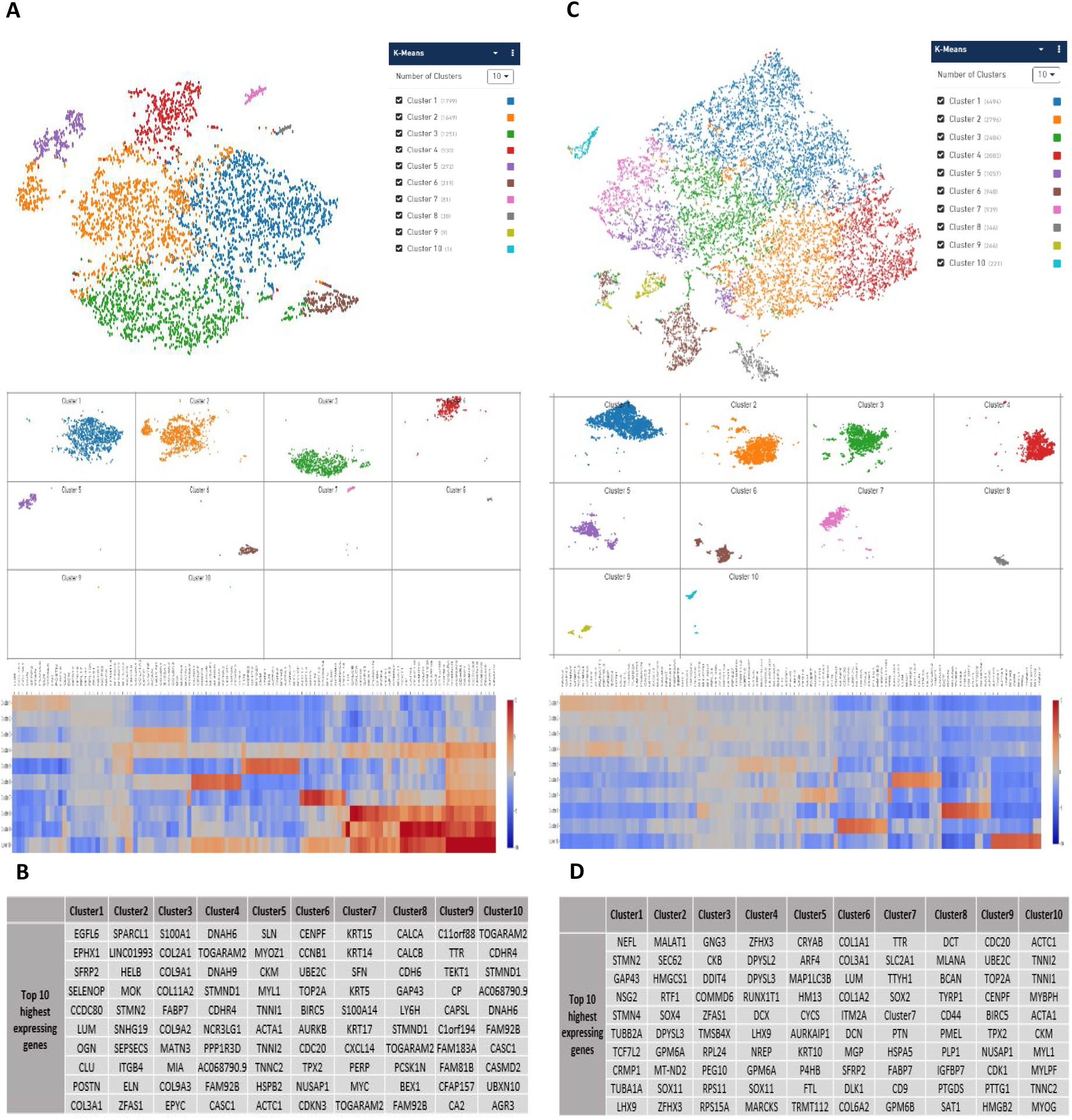
Single-cell RNA transcriptome clustering for ocular and cerebral organoids. (A, top) The t-Stochastic Neighbor Embedding (t-SNE) plot of 5849 cells from human ocular organoids derived from H1 human ES cells separated into 10 distinct clusters by K-means. Cluster 9 and 10 with low cell numbers were removed from subsequent AAV tropism analysis. The sequenced FASTQ files are processed by a modified Cell Ranger pipeline and visualized on the Cell Loupe software, the mean reads per cell is 122688 and the median genes per cell is 1022. (A, bottom) Each row represents the heat map of transcriptome expression profile of each of the 10 cell clusters that were separated by K-means. (B) A representative list of top 10 high-expressing genes for each cell cluster as used for identification of cell niche in the t-SNE plot. Refer to Supplementary Data I for full list of mRNA counts for each cluster of the ocular organoid. (C, top) The t-Stochastic Neighbor Embedding (t-SNE) plot of 15466 cells from human cerebral organoids derived from H1 human embryonic stem cells, separated into 10 distinct clusters by K-means. The sequenced FASTQ files are processed by a modified Cell Ranger pipeline and visualized on the Cell Loupe software, the mean reads per cell is 23315 and median genes per cell is 902. (C, bottom) Each row represents the heat map of transcriptome expression profile of each of the 10 cell clusters that were separated by K-means. (D) A representative list of the top 10 highly-expressing genes for each cell cluster for identification of the cell niche in the t-SNE plot. Refer to Supplementary Data II for full list of mRNA counts for each cluster of the cerebral organoid.

**Supplementary Figure 4.**
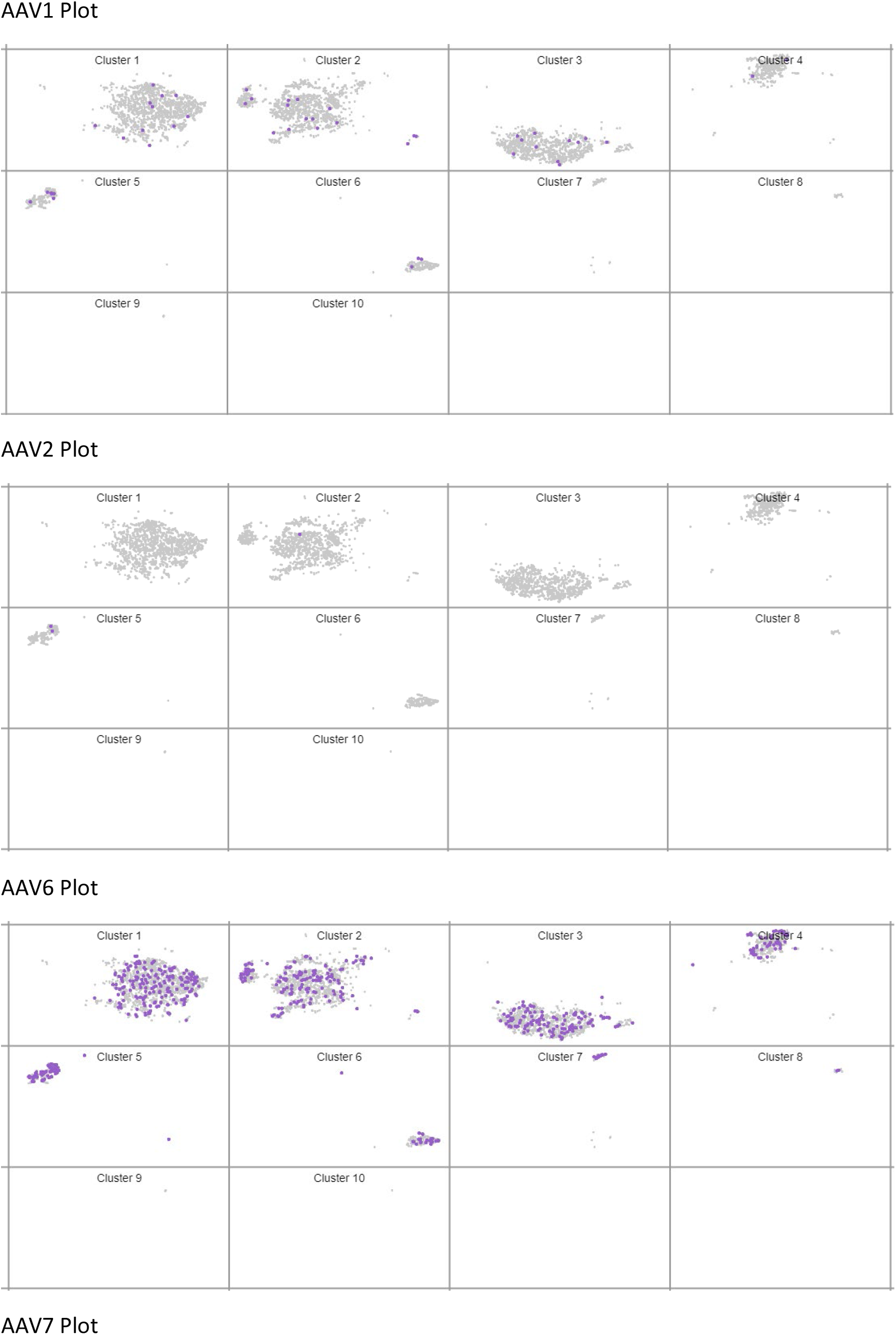
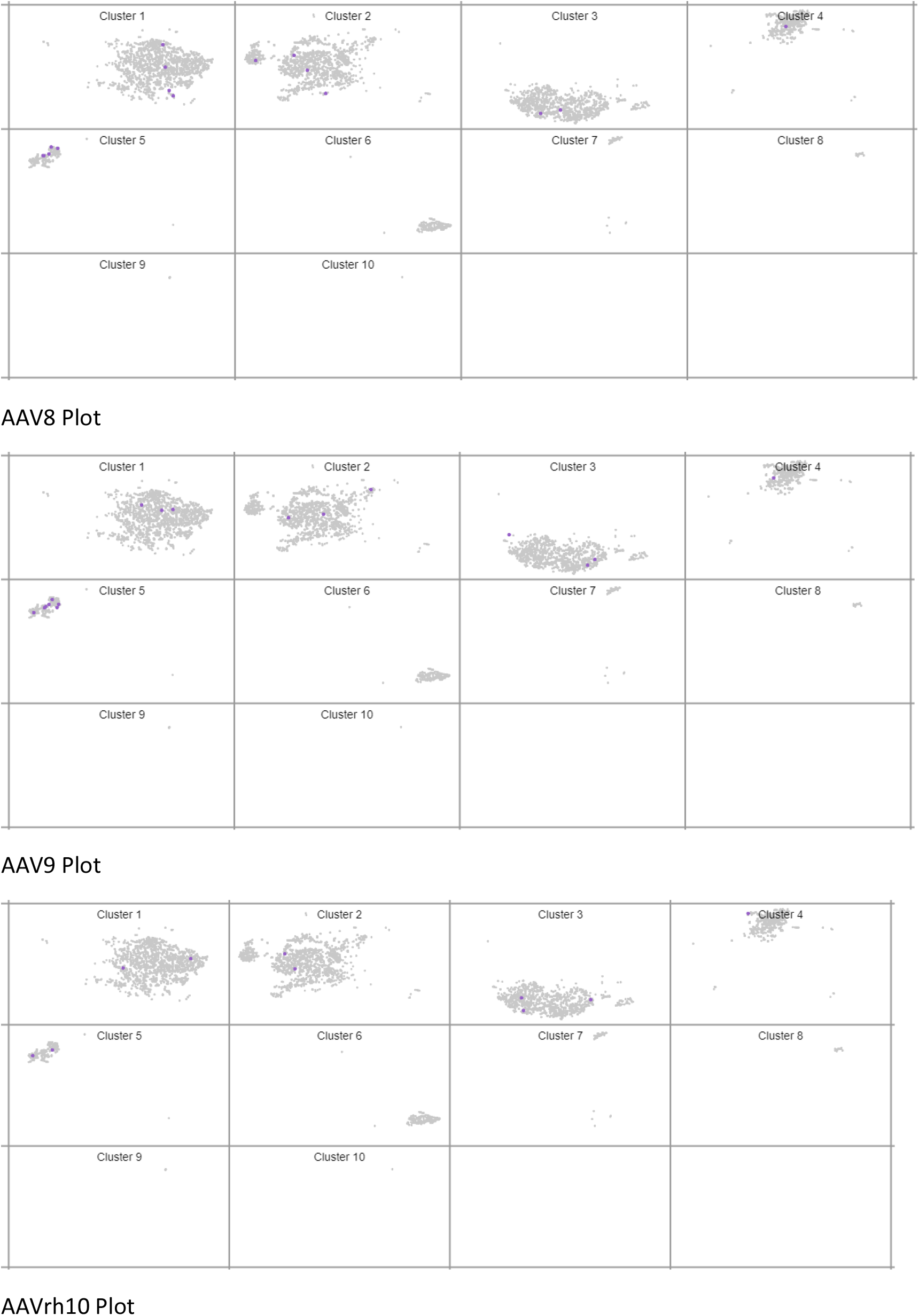
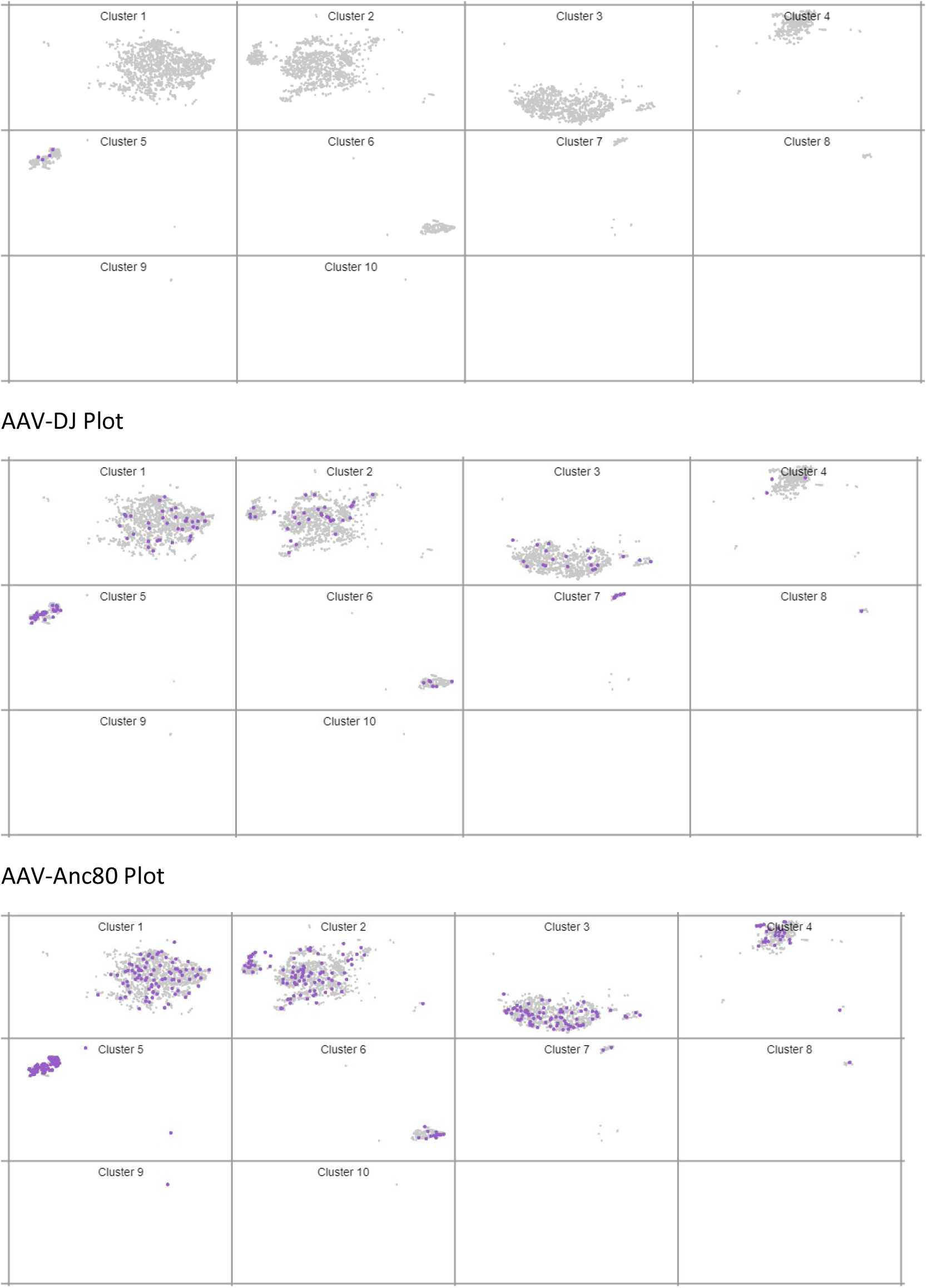
t-SNE plots showing individual cells transduced with different AAV serotypes (each serotype represented by one plot), in each of the 10 clusters of the ocular organoid.

**Supplementary Figure 5.**
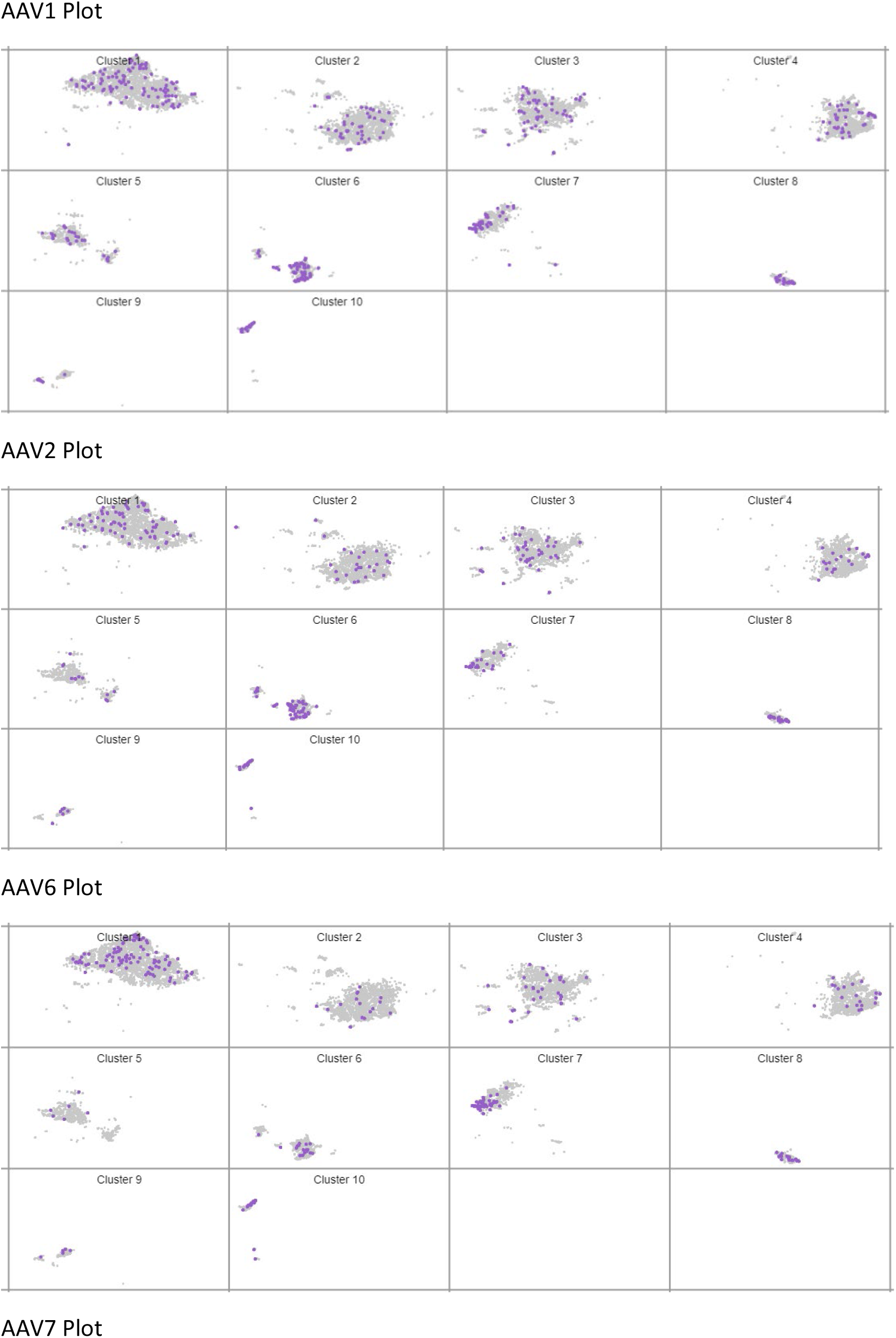
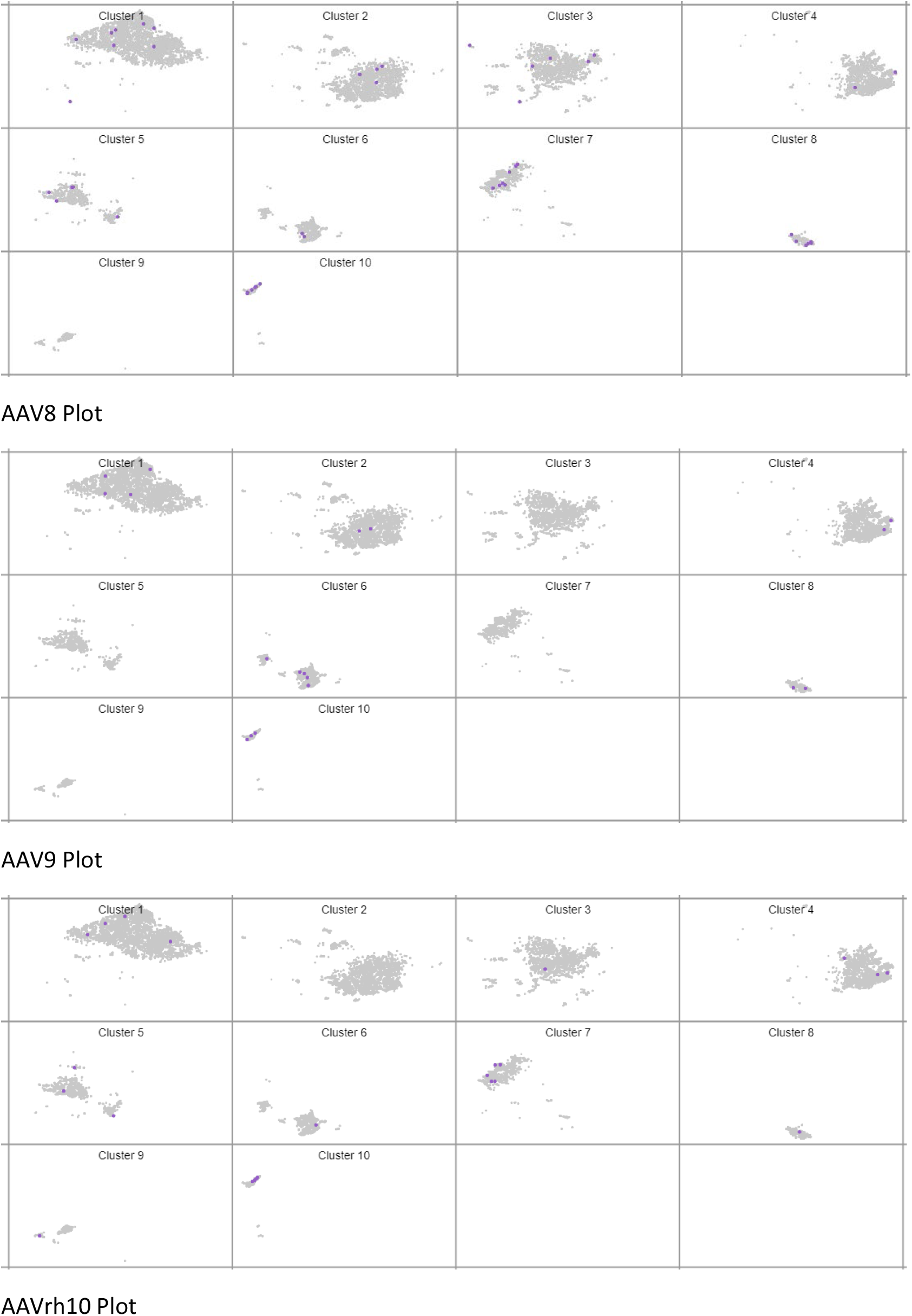
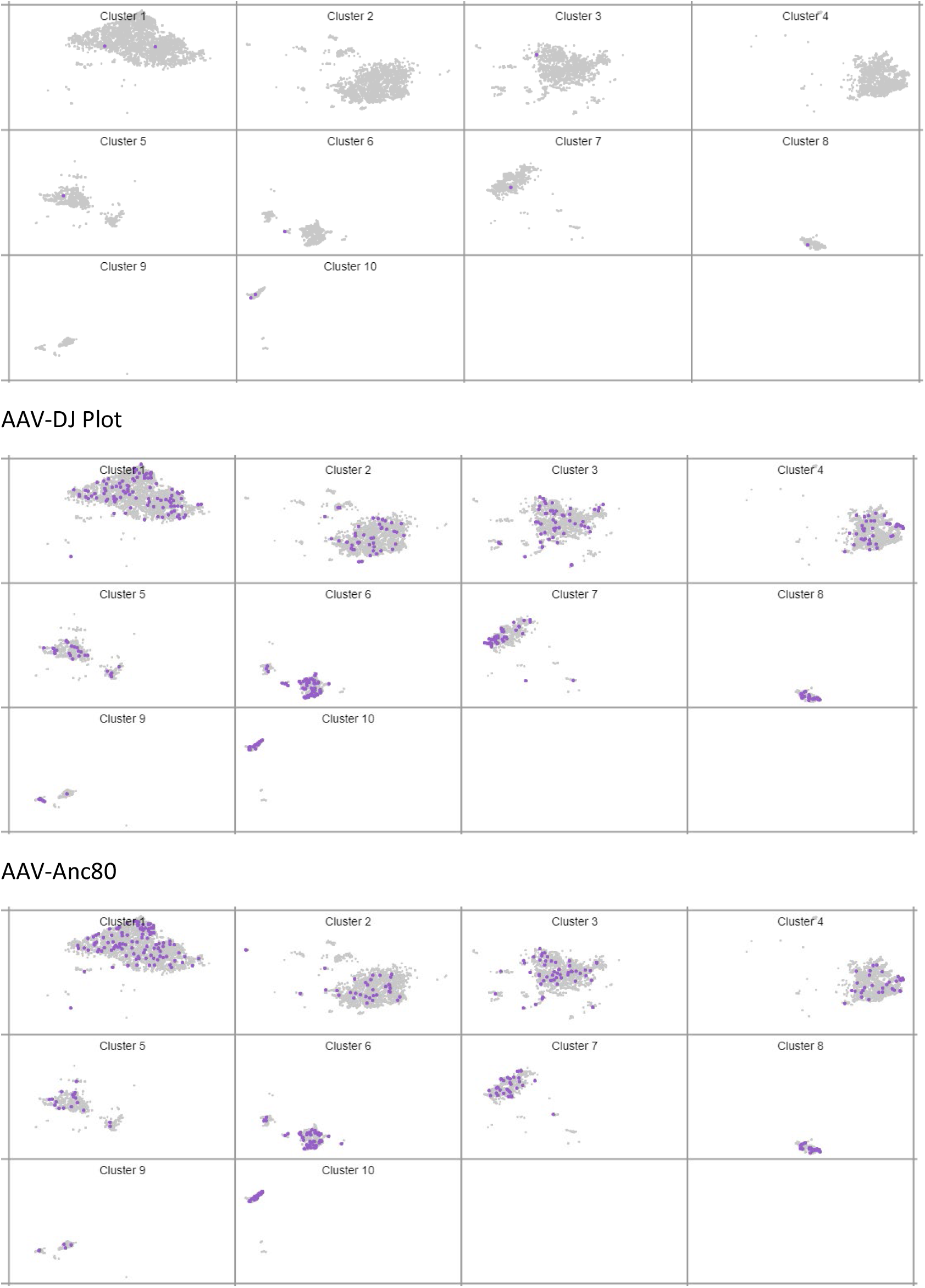
t-SNE plots showing individual cells transduced with different AAV serotypes (each serotype represented by one plot), in each of the 10 clusters of the cerebral organoid.

## Supplementary Data IV

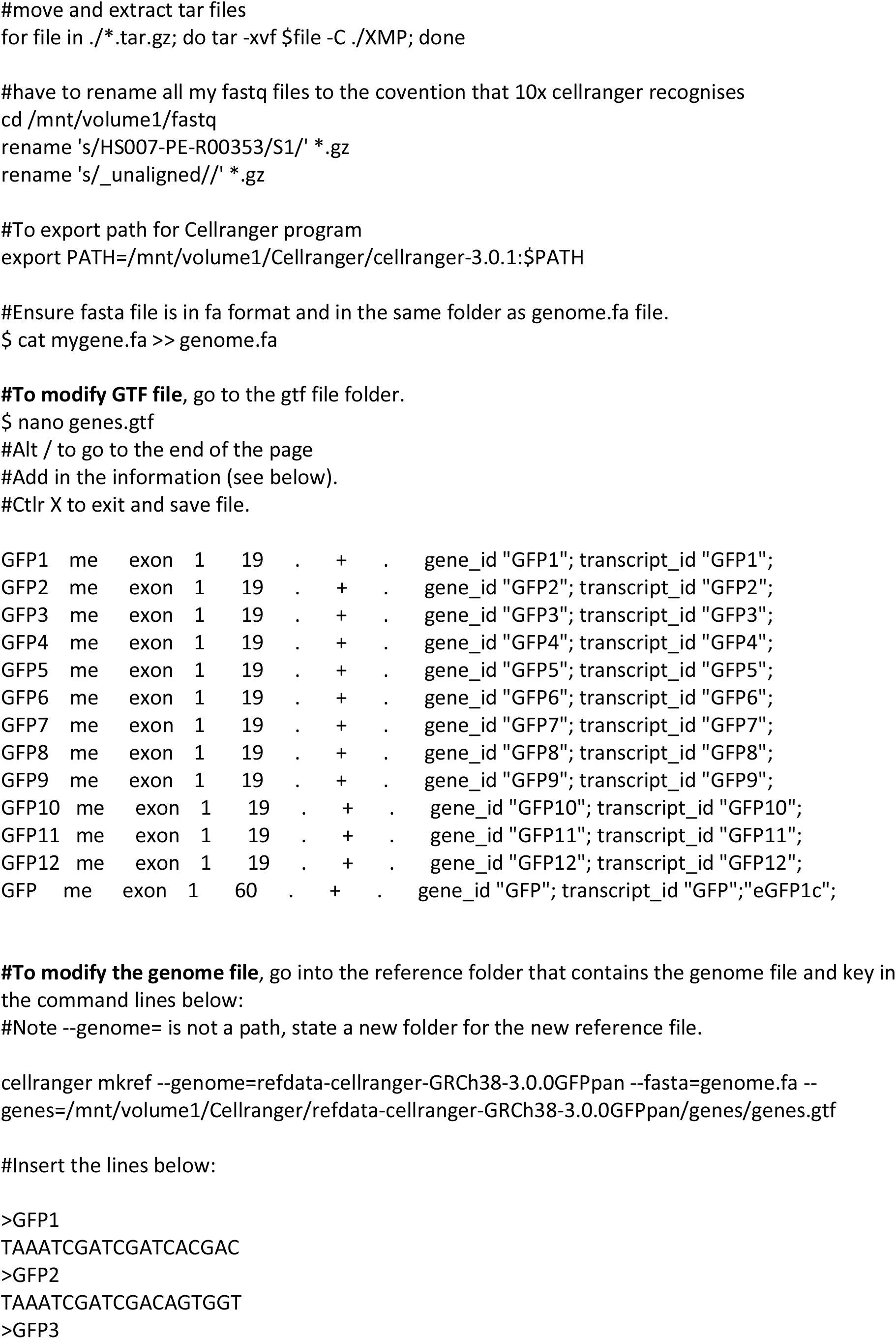
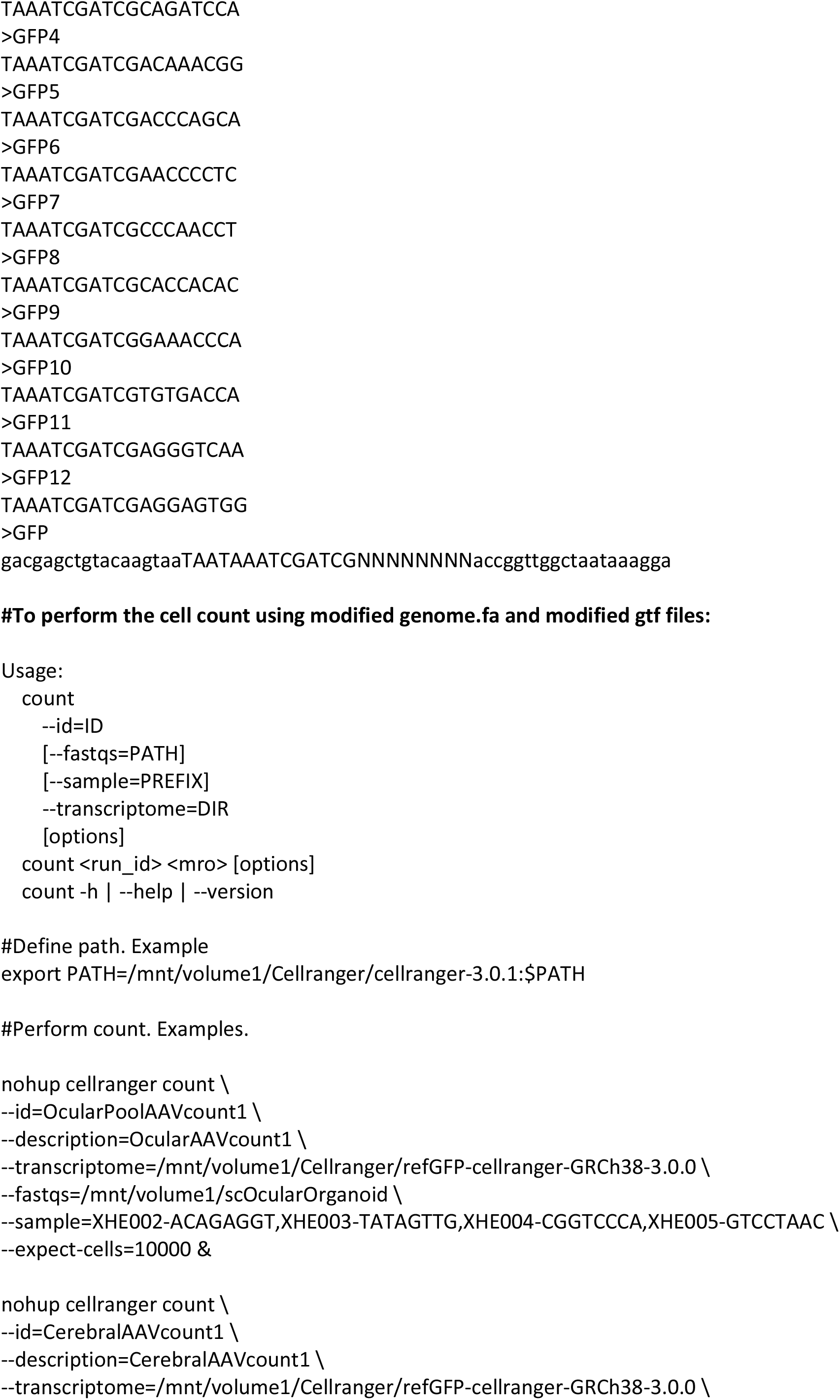
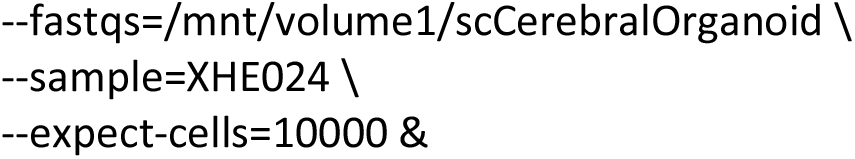

